# A microfluidic device for controlled exposure of transgenic *Ciona intestinalis* larvae to chemical stimuli demonstrates they can respond to carbon dioxide

**DOI:** 10.1101/2022.08.15.492342

**Authors:** Guillaume Poncelet, Lucia Parolini, Sebastian M Shimeld

## Abstract

The larva of the ascidian *Ciona intestinalis* controls a small repertoire of behaviours with a simple nervous system in which each cell is identifiable. As such it offers the prospect of building a cohesive cell-level picture of how a nervous system integrates sensory inputs to produce specific behavioural outcomes. Here, we report the development of a microfluidic chip in which larvae can be immobilised and exposed to chemical stimuli. We generate transgenic larvae in which the calcium ion reporter GCaMP6m is expressed in a defined population of cells, allowing us to record real-time neural activity following stimulation. We then use this to establish that some cell populations can sense dissolved carbon dioxide. We also leverage genome and transcriptome data coupled with molecular evolutionary analysis to identify putative chemoreceptors of the MS4A family in *Ciona*. Our study demonstrates that *Ciona* larvae can respond to dissolved carbon dioxide, identifies the cells that are likely responsible for chemosensation, and establishes a chip based imaging platform coupled with transgenic technology that could be adapted to establish where other stimuli are sensed and how such incoming signals are processed in the brain to yield behavioural output.

## Introduction

Understanding how animals receive, process and respond to stimuli is a major challenge in biology, but one that recent innovations in live cell imaging are helping to resolve. Animals with relatively simple nervous systems offer the prospect of imaging any or all individual neurons in whole live organisms. The larva of the sea squirt *Ciona* is one such system. *Ciona* embryos undergo a deterministic developmental program resulting in larvae with a small and defined number of individually-identifiable nerve cells [1, 2]. The connectome of the *Ciona* larval nervous system has been described [3] and single cell sequencing is generating a deep descriptive understanding of cell transcriptional identities [4, 5]. *Ciona* is also easy to genetically manipulate, with well-established methodology allowing simultaneous generation of hundreds of transgenic animals [6].

*Ciona* larvae are highly motile and integrate environmental cues to identify an appropriate site for settlement and metamorphosis [7]). Experiments have demonstrated that larvae can respond to light and gravity, with these stimuli detected by sensory cells embedded in the anterior swelling of the central nervous system known as the sensory vesicle (Figure 1A, B; [8-10]). Larvae also possess a small number of sensory cells in the epidermis (Figure 1C; [2, 11]). These include neurons in the tail (DCENs, VCENs and BTNs: see Figure 1 legend), ciliated neurons dorsal to the sensory vesicle known as the anterior and posterior Apical Trunk Epidermal Neurons (aATENs and pATENs), neurons embedded in the three anterior projections known as palps (Palp Sensory Neurons; PSNs), and the Rostral Trunk Epidermal Neurons (RTENs) that link the palps to the sensory vesicle. The aATENs and PSNs have been proposed to be chemosensory neurons [12, 13]. The palps also contain another cell type, the Axial Columnar Cells (ACCs: Figure 1C), which have also been suggested to be a site of chemosensation [14]. This was based on the observation that they express a 7 transmembrane domain encoding gene coupled with the knowledge that vertebrate Olfactory Receptor (OR) proteins also have 7 transmembrane domains, though *Ciona* and vertebrate proteins are quite divergent [15].To date, however, chemosensory function has not been experimentally demonstrated for these or any other *Ciona* cells.

**Figure 1.**
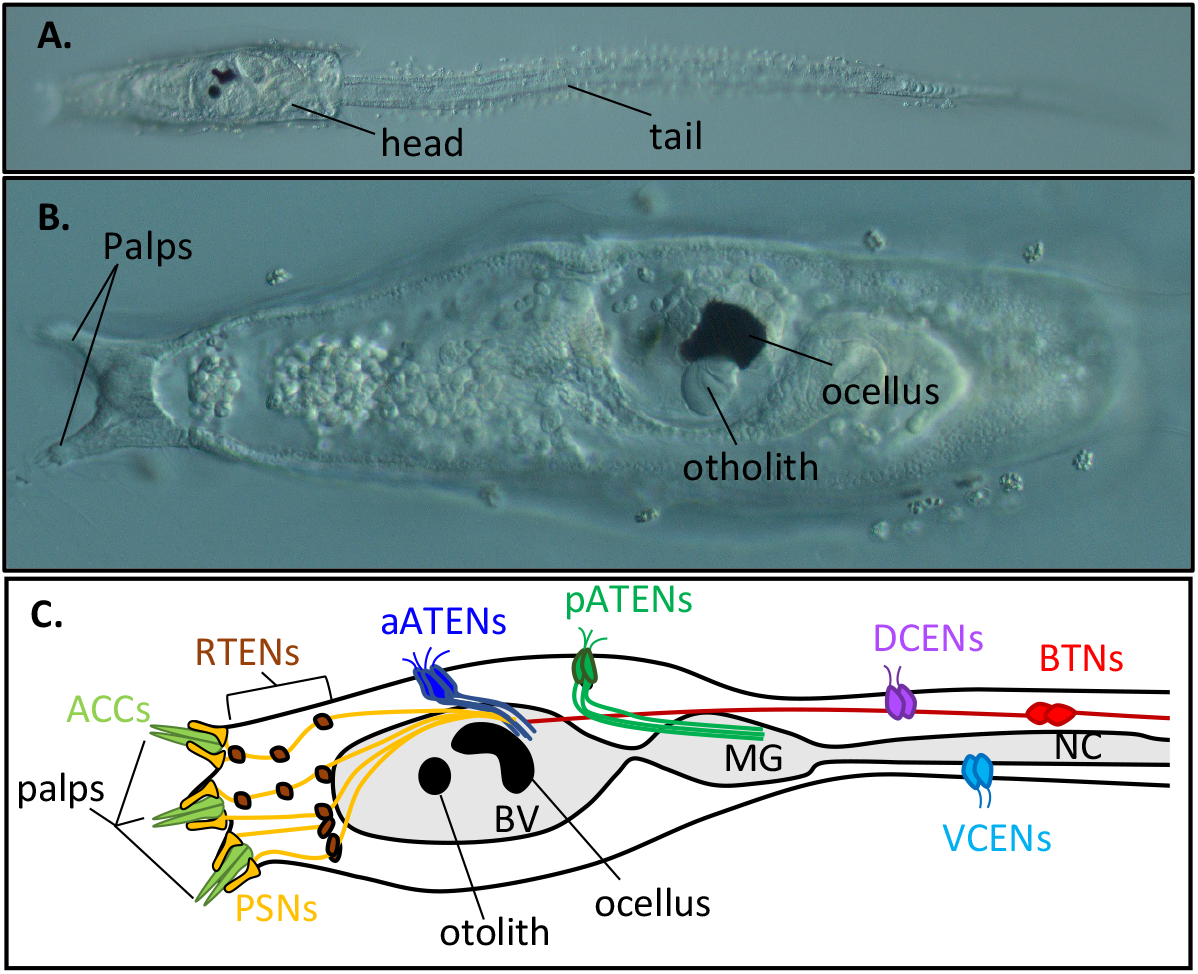
*Ciona* larva neural anatomy. (A) Motile *Ciona* larva at 22hpf. (B) Head of *Ciona* larva at 22hpf with palps, otolith and ocellus labelled. (C) Schematic drawing of *Ciona* larval head showing the various epidermal sensory cells: Bipolar Tail Neurons (BTNs), Dorsal and Ventral Caudal Epidermal Neurons (DCENs, VCENs), anterior and posterior Apical Trunk Epidermal Neurons (aATENs and pATENs), Rostral Trunk Epidermal Neurons (RTENs) andthe Palp Sensory Neurons (PSNs) and Axial Columnal Cells (ACCs).

Larval behaviour and ecology suggest *Ciona* larval chemosensation is likely to include sensation of biofilm-derived chemical cues for selecting an appropriate settlement site [16]. It may also include sensation of dissolved gasses such as carbon dioxide (CO_2_). The ability to detect CO_2_ is widespread in living-organisms and can help maintain homeostasis, locate food and avoid predators [17, 18]. The mechanism has not been well studied in most animals though some mammals can sense environmental CO_2_ through their olfactory systems using a type of olfactory sensory neuron (OSN) known as necklace OSNs. While most OSNs use OR genes to detect olfactants, necklace OSNs express a different complement of receptors and downstream signalling molecules [19].

One method of detecting neural activity is through Calcium ion (Ca^2+^) imaging using genetically encoded Ca^2+^ sensors such as GCaMP [20]. Previous studies have demonstrated GCaMP activity in *Ciona* following injection of *GCaMP* mRNA or electroporation of GCaMP-encoding plasmids [21-25]. In principle this technology should allow the imaging of individual sensory cells (and other neurons) while animals are exposed to defined stimuli, allowing sensory cell modality and downstream neural responses to be established. A major challenge, however, is the motility of the larvae: larvae swim vigorously making imaging of free-range larvae impossible. While anaesthetics can block this behaviour, doing so may compromise neural responses and hence also experimental outcomes. Other forms of immobilization are necessary if imaging is to link controlled sensory stimulation to neural and behavioural response.

Here we address chemosensation in *Ciona* from two angles. First we establish methodology for live Ca^2+^ imaging in *Ciona* larvae exposed to a controlled chemical stimulus: this includes the development of a microfluidic chip that allows *Ciona* larvae to be immobilized and imaged while exposed to potential olfactants, coupled with the deployment of a transgenic GCaMP system in which the reporter protein is expressed in defined cells. Second we mine *Ciona* genome and transcriptome resources to identify candidate receptor and other genes with the potential to be involved in chemosensation. Our results show that *Ciona* larvae can sense dissolved CO_2_ levels and identify a suite of genes expressed in palp cells likely to be responsible for this and other chemosensation.

## Materials and Methods

### Animals and electroporation

Adult *Ciona intestinalis* (*Ciona intestinalis* Type B) were collected from Northney Marina, Hayling Island, UK. They were maintained in a marine aquarium at 13°C and held under constant light to inhibit natural spawning. Gametes were removed by dissection and fertilization performed *in vitro*. Within 15 minutes of fertilization zygotes had their chorion and follicle cells removed [26]. They were then electroporated with plasmid DNA [27]. Transgenic embryos were raised on agarose coated petri dishes in filtered sea water until they reached the desired stage for the experiment, i.e. 20 hpf. All experiments were conducted on *Ciona intestinalis* (*Ciona intestinalis* type B) which is hereafter referred to as *Ciona*. Some genome and transcriptome data from two other *Ciona* species, *Ciona intestinalis* Type A (also called *Ciona robusta*) and *Ciona savignyi*, were also used: where this has occurred the full species is indicated.

### *DMRT>GCaMP6m* construct design and validation

The *DMRT* regulatory region was amplified from the *DMRT>GFP* plasmid [28] using Phusion polymerase (New England Biolabs) with primers: 5’-ACGAGATCTTAGTAGGGTGGAGGAAGATGG-3’ and 5’-ACGCCTAGGGCCAGTTA-AACGAACTGTTTG-3’. The *DMRT* regulatory region was inserted into the plasmid pAAV-hSyn1-mRuby2-GSG-P2A-GCaMP6m-WPRE-pA (Addgene) cut with AvrII and BglII restriction enzymes.

To validate GCaMP6m activity in *Ciona*, larvae transgenic for *DMRT>GCaMP6m* were immobilised in low-melting point agarose inside a perfusion slide (µ-Slide Luer, ibidi) perfused with sea water. GCaMP6m was excited at 488 nm and fluorescence detected with a Fluorescence Axiozoom V.16 microscope, equipped with a Plan-NEOFLUAR Z 2.3x (NA 0.57). GCaMP6m fluorescence was recorded for 30 seconds (300 frames at 10 frames per second). The sea water solution was then switched, and the slide was perfused with 1M calcium ionophore A23187 diluted in sea water. GCaMP6m fluorescence was then also recorded for 30 seconds (300 frames at 10 frames per second). Identification of the responding cells was undertaken using Fiji by calculating the standard deviation through all 300 frames in both conditions.

### Microfluidic chip fabrication

Standard soft lithography was used to fabricate the mould [29]. The photomask was designed with AutoCAD (2019 free student version, Autodesk, Inc.). Premium grade high resolution film photomasks (features down to 5μm) were printed by Micro Lithography Services Limited, Chelmsford, UK. The photomask source file is available as .dwg file (Additional File 6). The chip mould with a uniform height of 23.5μm was obtained by spin-coating a silicon wafer (4 inches; Siltronix, France) with a negative photoresist (SU-8 2035, MicroChem Corp., Newton, MA, USA) according to manufacturer’s instructions. Devices were produced by pouring onto a mould a prepolymer mixture of polydimethylsiloxane (PDMS, Sylgard 184 silicone elastomer kit, Dow Corning Corp.) with a 1:10 ratio of curing agent and curing at 65°C for a minimum of 4 hours. PDMS blocks were then irreversibly bound to a 1mm thick standard microscope slide (25mm x 75mm) by a 1 minute treatment in a plasma oven.

### Equipment organisation, chip loading, imaging and recording

Equipment organisation is shown in Additional File 1. A customised 3D-printed chip holder was used to maintain the chip under an upright microscope. The chip holder source file is available as .stfl file (Additional File 7). Inlet pressure was controlled using an MFCS™-EZ (Fluigent) having an integrated positive pressure source with three pressure channels with ranges from 0-345 mbar. This was connected to a computer running the Fluigent Microfluidic Automation Tool which controlled flow rates in each channel. Each channel drew from a separate tube filled with a test solution. The MFCS™-EZ was positioned next to the microscope stage, with a constant tubing length of 30cm used to connect tubes and chip. The electroporated transgenic larvae were introduced in the chip by sucking them up into a PFT tube plugged to a metallic needle (Mircolance #20, 302200, BD, USA) connected to a plastic syringe (Luer Plastipak, BD, USA). The animals inside the tube were manually fed inside the introduction inlet by applying a light pressure on the plastic syringe to slowly push the larvae inside the trapping channel.

Calibration recordings were done with a Carl Zeiss Axioskop 2 plus microscope mounted with a Zeiss AxioCam HRc camera. Fast Green FCF (E143) was dissolved at 0.5gmL^-1^ in the two side streams to visualize their moving boundaries (Additional File 2, Additional Files 12-14). A single frame was taken every 6.8 milliseconds. The mean grey value through all frames was calculated using Fiji, in a square region of interest (ROI), constant in size and position, located in the trapping channel. Minimal and maximal pixel values, which according to the Beer– Lambert law corresponded, respectively, to the absence of Fast Green FCF and its maximum concentration, were normalized between 0 and 1 (Additional File 2). Measurements were made successively 60 times and these data superimposed to quantify variability of the stimulus timing with cycles and stimulus sides (Additional File 2). This revealed that the onset took 390 milliseconds (msec) on average with a duration variability of 197 msec. The offset took around 27 msec, with no detectable variability in duration. The experiment demonstrated the reproducibility of the stimulation onset/offset. Once these conditions for boundary control were established, the dye was no longer used. Calcium recordings were conducted on 20-24 hours post fertilisation (hpf) larvae at room temperature.

### Preparation of stimulus test solution

Artificial seawater (ASW) with a salinity level of 35 ppt, a carbonate hardness (dKH) of 8 and a pH of 8 was obtained by mixing 35g of pharmaceutical grade sea salt (PRO-REEF Sea Salt) per litre of ultrapure water. CO_2_ was dissolved in ASW using a SodaStream with a built-in CO_2_ cylinder. The made-up ASW solution with infused CO_2_ (ASW-CO_2_) had a lowered pH of 6. This decrease in pH relates to an estimated CO_2_ concentration of 129.3mgL^-1^ (ppm) in comparison to non-injected ASW at 1.1mgL^-1^ (ppm) of CO_2_, based on calculations from [30]. Due to the lowering effect of CO_2_ on the pH, a control ASW solution was made with a corrected pH of 6 by gradual addition of 1M HCl. The stock solutions were kept at 18°C in sealed glass bottles for long-term storage and loaded in 15mL Falcon tubes on the day of the experiment to avoid possible detection of dissolved substances from plastic containers.

### Image analysis

GCaMP6m fluorescence excited at 488 nm wavelength was detected with a Fluorescence Axiozoom V.16 microscope, equipped with a Plan-NEOFLUAR Z 2.3x (NA 0.57). 3000 images were taken at 10 fps (100ms exposure time) in 300 seconds. A pixel binning of 5×5 was done to increase signal-to-noise ratio. Movement artefacts on the raw calcium recordings were first corrected in FIJI (v.2.1.0) using the plugin TurboReg with rigid body transformations [31]. Mean fluorescence intensity was then calculated from ROIs drawn manually in FIJI. The attribution of a ROI to a specific embryonic region relied on its relative position guided by anatomical landmark recognition such as the palps, tail, ocellus and otolith. Further data analysis was done in Excel, following the guidelines of Akira Muto (http://akiramuto.net/archives/148). Traces were plotted as ΔF/F_0_=(F_t_-F_0_)/F_0_, with F_t_ the fluorescence intensity at time t, and F_0_ the mean fluorescence value over a 5s time window during the initial 30s resting state. The fluorescence change (ΔF/F_0_) was calculated after subtracting the background fluorescence. The raw recordings are available as .avi files (Online resource 4).

### Immunofluorescence staining of larvae

Transgenic swimming larvae (22-24hpf) generated by electroporation were fixed with 4% paraformaldehyde (PFA) in 1× phosphate buffered saline (PBS) for 30 min, washed three times with PBS, then gradually dehydrated to 100% methanol and stored at −20 °C. After stepwise rehydration to 100% PBS, samples were permeabilized with 0.2% (w/v) Triton X-100 in PBS (PBS-T). Non-specific antibody binding was blocked with 20% (w/v) heat treated sheep serum (HTSS) in PBS-T for >1 h at room temperature and samples were then incubated with primary antibody in HTSS-PBS-T. Primary antibodies were: rabbit-anti-Ci-ßγ-Crystallin [32], mouse-anti-mCherry monoclonal (Antibodies.com, A104343) all diluted 1:500 at 4 °C overnight. After several washing-steps with PBS-T at room temperature, they were incubated in secondary antibody, either goat anti-rabbit Alexafluor 488 (Molecular Probes, A11008) or goat anti-mouse HRP conjugate diluted 1:500 in BSA-PBS-T. After 5 washes in PBS-T, a tyramide signal amplification (TSA) was performed by addition of tyramide solution made of TSA buffer (2M NaCl, 100mM Borate buffer pH 8.5), 0.5% H_2_O_2_, 0.1% 4-Iodophenylboronic acid (4IBPA) and 0.2 % 5-TAMRA fluorescent dye (Abcam, ab145438) for 30 min. Embryos were washed 5 times in PBS-T for 30 min and once overnight. Nuclear DNA was stained with DAPI (1:1000 dilution). DAPI, Alexa 488 and 5-TAMRA fluorescence were excited at 405, 488 and 559 nm, respectively, and recorded with an Olympus Fluoview FV1000 confocal microscope equipped with 60X and 100X oil-immersion objectives. Samples were washed with PBS several times, mounted with 50% (w/v) Glycerol in PBS on a coverglass-bottom petri-dish. Stacks of optical sections from 0.3 μm to 0.8 μm were acquired sequentially (one channel at a time) and z-projected. Images were analysed with Fiji (Version 2.0.0).

### Analysis of Single-cell RNAseq data

Processed scRNAseq data from *Ciona intestinalis* Type A (*Ciona robusta*) previously published by [5] and re-analyzed by [14] using Seurat v3 [33] identified a cluster of palp ACCs according to the highest relative expression of βγ-Crystallin [32]. Differential gene expression analysis identified genes with transcripts enriched in the ACCs compared to neural cells. The full code for this is available at https://osf.io/5dc4u / (see [14] for details).

### Amino acid conservation analysis of ascidian MS4A proteins

FASTA format sequences of the specified *Ciona intestinalis* Type A (*Ciona robusta*) and *Ciona savignyi* MS4A proteins were downloaded from the ANISEED database (http://www.aniseed.cnrs.fr/). A multiple alignment with a corresponding score for amino acid conservation was obtained using the PRALINE program with default settings where the least conserved amino acids were given a 0 score and the most conserved amino acids were assigned a 10 [34]. Topographical representations of the MS4A family member that was used on the top line of the alignment were made using the Protter program by manually entering the topographical orientation of the protein as predicted by TOPCONS.

### Diversifying selection analyses of ascidian *MS4A* genes

Ascidian MS4A amino acid and nucleotide coding sequences were retrieved from the Aniseed database (Additional File 8). When a gene had more than one predicted isoform, the sequence that contained the longest open-reading frame was selected. The consensus topology of each MS4A amino acid sequences was assessed with TOPCONS and only those with a characteristic tetraspanning topology were used [35]. A multiple alignment of amino acid sequences was done with Muscle using default parameters. Manual trimming of the alignment was done in AliView [36] to remove columns containing gaps in the majority of sequences. The N- and C-terminal domains predicted via TOPCONS were also removed because of uncertainty over alignment in those regions. Maximum likelihood phylogenetic trees were built in RAxML (GUI, v2.0.3) with the best substitution model suggested (CPREV+G+F) and 1000 bootstrap to support nodes. The final dataset consisted of 130 sequences and 100 aligned characters. Species-specific clades containing recently diverging paralogues were identified in the tree and considered when there were at least four sequences. In each clade selected, the complete coding sequences of the *MS4A* genes were re-aligned with Clustal Omega using default parameters and a corresponding codon alignment was obtained using PAL2NAL with all gaps removed [37]. The N- and C-terminal regions of the codon aligned *MS4A* sequences were too variable and hence not included for the subsequent statistical tests. Nucleotide-based phylogenetic trees were done with MEGAX using Kimura-2 parameter model to obtain branch lengths. The codon alignments and corresponding trees were fitted to codon-based models implemented in EasyCodeML to estimate non-synonymous to synonymous substitution rate (dN/dS) ratios (ω) across paralog sequences [38, 39]. These models are: M0 (one average ratio ω), M1a (neutral; codon values of ω fitted into two discrete site classes between 0 and 1), M2a (positive selection; like M1a but with one additional class allowing ω>1), M7 (neutral; value of ω following a β distribution with ω=1 maximum), M8 (positive selection; like M7 with one additional class allowing ω>1). These different models, all assume that ω ratio is the same across branches of the phylogeny but different among sites (codons or amino acids) in the alignment. Likelihood-ratio tests were used to compare the fit of these models to the sequence data. Support for positive selection was identified when M2a provided better fit than M1a or when M8 provided better fit than M7 [40]. The M1a-M2a comparison is the most stringent but can lack power to detect signatures of diversifying selection compared to the M7-M8 models comparison, which imposes less constraints on the distribution of ω, but may have a higher rate of false positives [41]. EasyCodeML was run in preset mode using default settings. The Bayes empirical Bayes (BEB) procedure executed in EasyCodeML was used to identify amino acid residues evolving under positive selection when the likelihood-ratio test had a significant result for any of the pairwise comparisons of codon models. The standard threshold for determining amino acid sites under selection is a posterior probability of 0.95 [42]. The predicted sites under positive selection were mapped across the predicted MS4A protein domains using the PROTTER program by manually entering the topographical orientation of the protein predicted via TOPCONS. To identify sites that have experienced purifying selection (p-value ≤ 0.10), the Fixed Effects Likelihood (FEL) method present in the HyPhy package was used [43].

## Results

### Construct design and verification

We first sought proof of principle for Ca^2+^ detection with the genetically-encoded calcium sensor GCaMP6m in live *Ciona* larval heads. We selected a GCaMP6 because *GCAMP6s* mRNA has been used successfully by injection in *Ciona* [22] and *Platynereis* [44]. We specifically selected Gcamp6m because it has a faster kinetic response [20]. We targeted GCaMP6m to a defined part of the head using the *Ciona DMRT* regulatory region [28] which expresses in rows III-VI of the neural plate at the mid-gastrula stage. Rows III and IV contribute to the CNS, including the anterior sensory vesicle, while rows V and VI contribute to the palps, oral siphon primordium and peripheral nervous system [45, 46]. To determine whether this construct was correctly expressed and could be seen and recorded responding to cytosolic Ca^2+^change, *Ciona* larvae transgenic for *DMRT>GCaMP6m* were immobilised in low-melting-point agarose in a perfusion slide and fluorescence recorded before and after the addition of the calcium ionophore A23187. Responding cells were identified by calculating the standard deviation through all frames to identify the pixels whose value fluctuated the most (Figure 2).This experiment demonstrated that the GCaMP6m could be delivered to appropriate cells by transgenesis, and that its fluorescence could be recorded in live animals.

**Figure 2.**
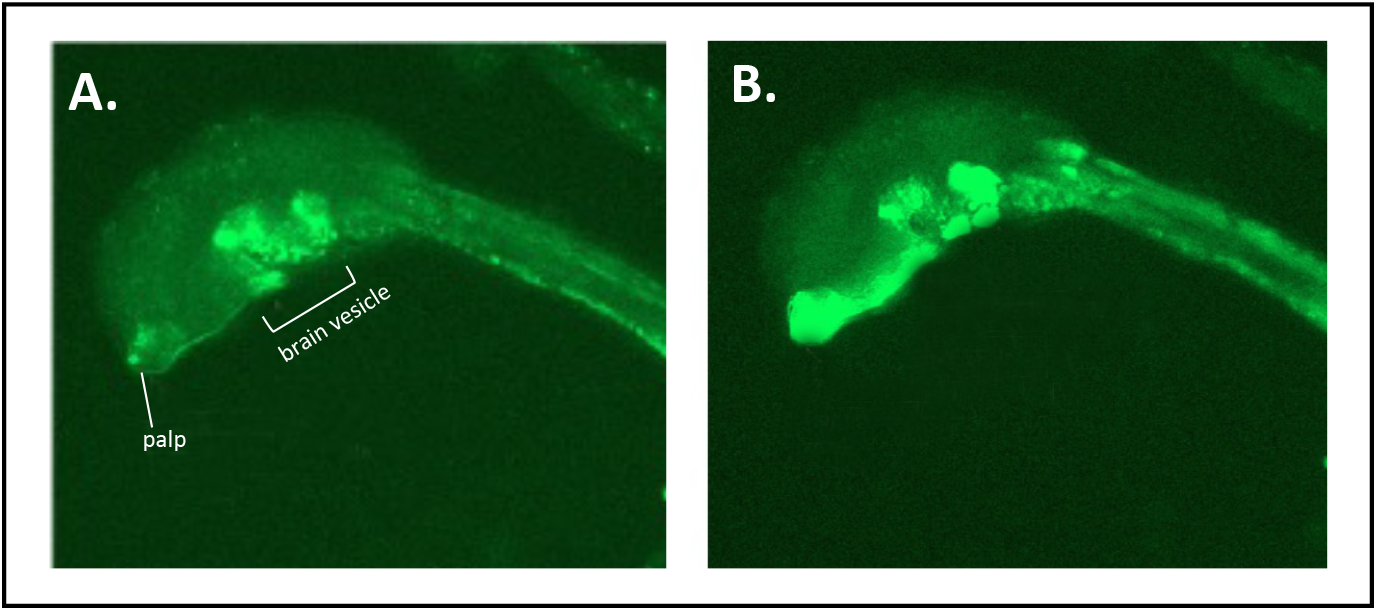
Validation of GCaMP6m fluorescence in transgenic *Ciona*. (A) *Ciona* larva perfused with sea water or (B) sea water with 1M Ca^2+^ ionophore A23187.

### Chip design and validation

To our knowledge, no prior study had attempted to introduce a *Ciona* larva into a microfluidic chip. Therefore, the first step was to identify the relevant dimensions to trap live *Ciona* larvae. In *Ciona* the larval head is the widest part of the body and measurements of 50 larvae revealed that the maximum head width was about 89μm (with a standard deviation of 10μm, 95% confidence interval bounds 82.15 to 95.7μm). We first trialled a modification of the chip design developed for immobilisation and imaging of larvae of the polychaete *Platynereis* [44]. This modified chip consisted of a trapping channel with a constant height of 60μm, with a width that decreased linearly along its main axis from 150μm to 80μm. While this design proved suitable for loading and initial containment of larvae, flow pressure caused larvae to be pulled back into the introduction channel or pulled beyond the trapping zone. This is because the heads of *Ciona* larvae can be deformed by pressure and lack the chaetae of annelid larvae which may help them lodge in position.

To circumvent these limitations, we trialled a variety of chip designs before settling on a symmetrical chip organised around a central trapping channel of width 500μm, length 2cm and height 23.5μm, which holds the animals dorso-ventrally (Figure 3A, Additional File 6). An animal introduction inlet is situated in the middle of the trapping channel. The chip includes three inlet channels which deliver the test solutions into a chamber upstream of the trapping channel, which has a width of 2mm. This splits into the trapping channel plus two lateral channels with a width of 1mm each. These come back together at the back of the chip where one outlet channel is sited.

**Figure 3.**
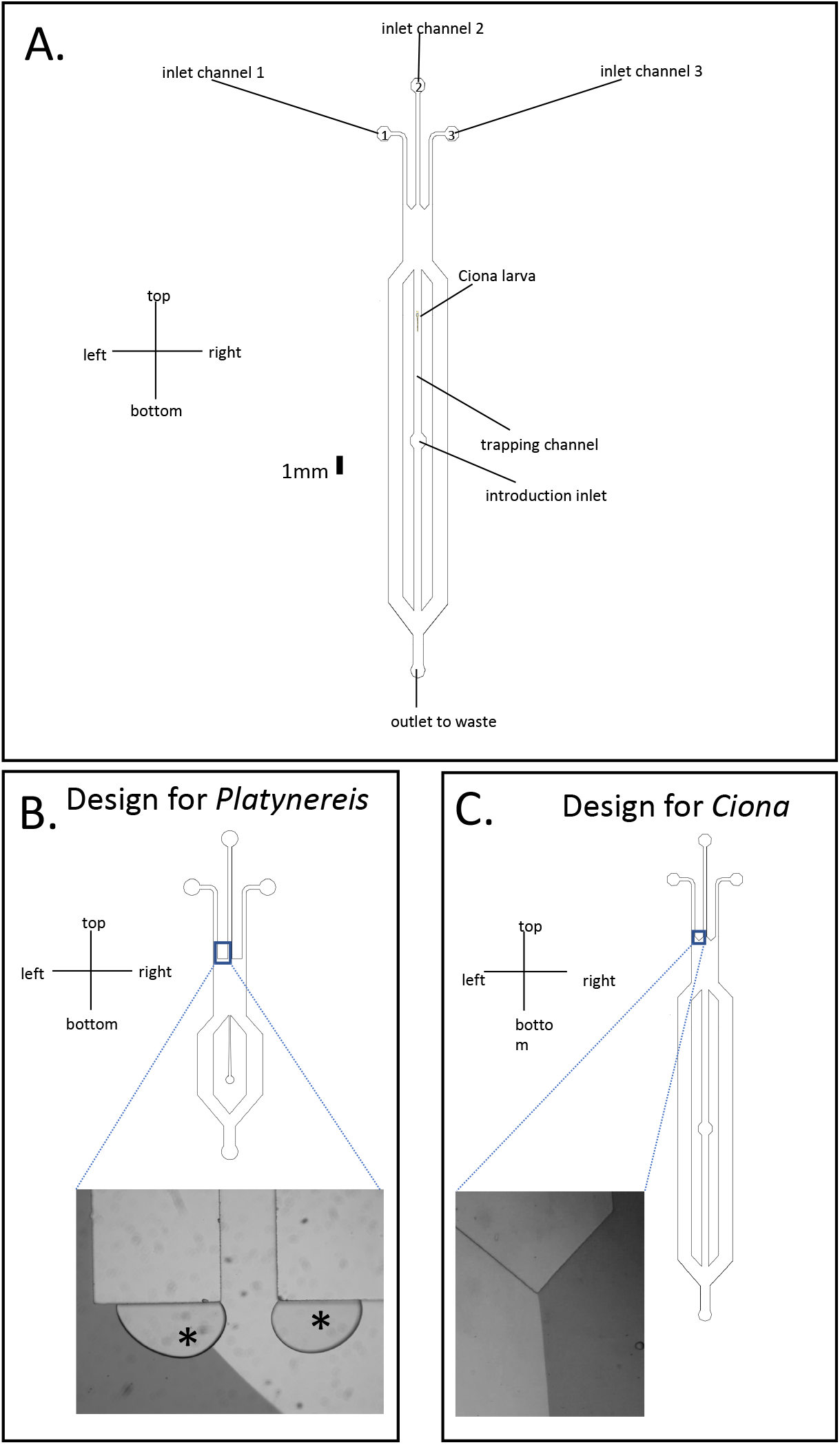
(A) Microfluidic chip design used in this study. Scale: 1 mm. (B) The chip design of [44] for *Platynereis* traps air bubbles visible on the inserted photograph (black asterisk) on flat surfaces located before the main chamber. (C) In the modified design used in this study the surfaces are triangular instead of flat precluding bubble retention. In B and C fluid flow is from top to bottom in the pictures, and flow boundary is visible as one stream is labelled with fast green (dark grey) and the other is unlabelled water (light grey). Design file is available as Additional File 6

This chip allows the introduction, immobilisation and recording of multiple animals at the same time. We also noted that aspects of the chip design of Chartier et al (2018) were enhancing the trapping of air bubbles, which may affect fluid flow behaviour (Figure 3B). Bubbles can be removed by degassing systems, however, this was not desirable here as the stimulus to be tested was also a gas. This issue was resolved by altering the chip design to make some surfaces triangular instead of flat (Figure 3C). This did not affect the smooth transition of flow boundaries but enabled bubbles trapped in the chip to be flushed out.

### Establishing laminar flow and testing the response to CO_2_

We first established flow conditions for exposing larvae to stimulus or control in a defined time frame. Fast Green added to the two lateral inlets allowed them to be visualised. We found maintaining a total pressure of 200mbar in the chip at all times, with the central inlet at 80mbar, allowed the controlled transition between exposure to central or lateral streams (Figure 4: a specimen recording is shown in Additional File 12). In experiments one side stream contained a high CO_2_ stimulus solution (S) and the other a pH-matched control (C) of sea water with a corrected pH value equivalent to that of the high CO_2_ solution.

**Figure 4.**
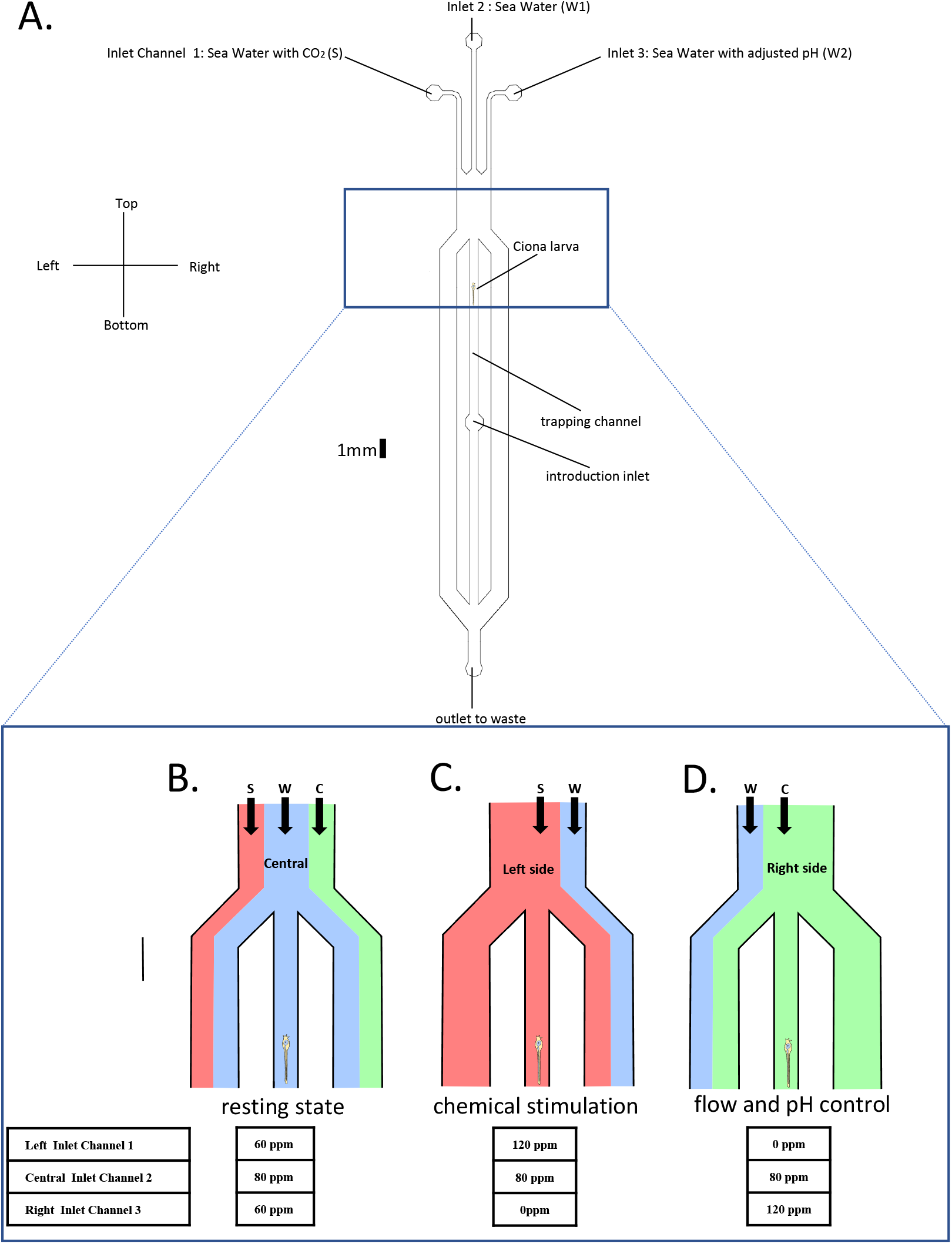
Experimental set-up for exposing Ciona larvae to CO2 enriched sea water. (A) Microfluidic chip design with inlets labelled to indicate solution flow. (B-D) Schematics of the different flow patterns used in the experimental protocol. Flow direction is from top to bottom. (B) In the resting state, the larva is exposed to a continuous flow of artificial sea water (W, in blue) while the stimulus (S, in red) and the flow control (C, in green) are flowing on the side streams. (C) During chemical stimulation, the flow boundaries move to hit the larva with the stimulus. (D) In flow control the larva is instead hit with the control pH-matched stream. The pressure value for each inlet used to obtain the specific flow patterns is indicated on the bottom. Scale on A is 1 mm.

In total seven larval heads were imaged (Figure 5, Additional File 3). Parts of the head were designated as Regions Of Interest (ROIs). To define these we first examined recordings by eye in Fiji, looking for regions which showed obvious and rapid changes in fluorescence. In most larvae more than one such ROI could be identified, though sometimes this was not possible as the transgene showed some mosaicism, and sometimes larvae were damaged during loading (Additional File 3). We were not able to identify individual cells within ROIs as the *DMRT* enhancer expresses quite broadly, however their positioning relative to anatomical landmarks suggested their identity.

**Figure 5.**
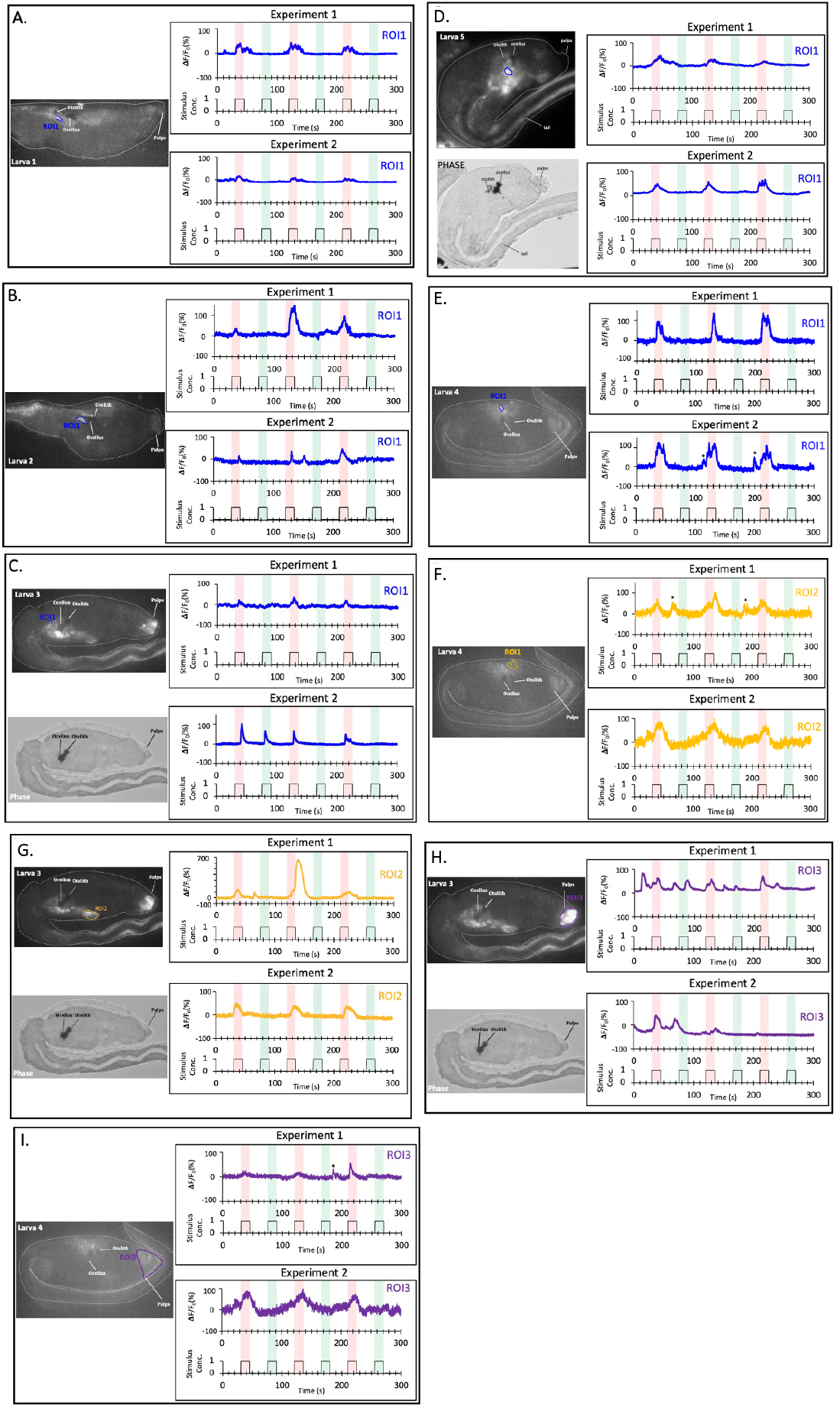
Experimental responses are registered by calcium imaging in Dmrt+ cells. The fluorescence image shows the region of interests (ROI) used to make the calcium activity traces. The graphs show the temporal patterns of fluorescence intensity for each ROI. The red bar corresponds to exposure to the solution with high CO2. The green bar is exposure to pH control. Asterisks mark random calcium peaks happening outside the zone of chemical exposure (red). Note E, F and I are separate ROIs recorded for the same larva, as are C, G andH.

A small population of cells near the ocellus and otolith were found to synchronise perfectly with the flow protocol, with strong reporter activity coinciding with the CO_2_ stimulus but not with the pH matched control (Figure 5A-F). From their position these could be aATEN epidermal sensory neurons (Figure 1C). However, sensory vesicle neurons also lie under the aATENs in this area and we were not able to conclusively distinguish between these cell populations. Responsive cells were also seen anterior to the brain vesicle (Figure 5G), positioned between the palps and brain vesicle. This corresponds to the position of the RTENs. These neurons connect the palps to the sensory vesicle, so if palp cells were involved in detecting stimuli we would expect this signal to pass via the RTENs. Finally, when a region of interest was drawn in the palp area (Figure 5H,I), reporter spikes were registered when stimulated by both increased CO_2_ solution and the pH-matched control solution.

### Genomic insights into the molecular basis of CO_2_ sensing in *Ciona*

Previous studies seeking insight from genomic data into chemosensation in *Ciona* have focused on OR genes or their invertebrate equivalents [47-52]. These studies have failed to demonstrate clear orthologues of either gene group, though some divergent GPCRs were identified. CO_2_ sensing has not been studied in *Ciona* and remains poorly understood in most animals, though has been traced to the necklace OSNs in mammals where some of the mechanism is known. We mined previously published [14] single cell RNA-seq data, finding orthologues of genes involved in mammalian CO_2_ sensing were enriched in the palp ACC cells (Table 1). These include genes encoding carbonic anhydrase CAII which catalyses the hydration of CO_2_ to H_2_CO_3_, the receptor guanylate cyclase GUCY2D which converts bound GTP to cGMP, the cyclic nucleotide gated channel CNGA3 to which cGMP then binds and PDE2A converts the cGMP into GMP. scSeq data were not, however, resolved to the level where ATENs or RTENs could be investigated.

**Table 1.** Selected genes with transcripts enriched in the ACCs.

| ANISEED ID | KH ID | Top BLASTP hit in human | AveLog(FC) | p value | Adj. p value | Pct-1 | Pct-2 |
| --- | --- | --- | --- | --- | --- | --- | --- |
| Cirobu.g00004428 | KH.C2.249 | CNGA1;CNGA2; CNGA3 | 0.705277326 | 2.34E-43 | 3.15E-39 | 0.31 | 0.023 |
| Cirobu.g00000663 | KH.C1.423 | CA1; CA2; CA7 | 0.260192973 | 5.76E-05 | 0.77389751 | 0.131 | 0.058 |
| Cirobu.g00010171 | KH.C9.608 | PDE2A; PDE8A; PDE9A | 0.664092861 | 1.21E-15 | 1.62E-11 | 0.225 | 0.037 |
| Cirobu.g00003495 | KH.C13.90 | GUCY2D; NPR1; NPR2 | 0.626634328 | 3.26E-19 | 4.38E-15 | 0.259 | 0.105 |
| Cirobu.g00000727 | KH.C1.484 | / | 0.385613806 | 2.26E-11 | 3.03E-07 | 0.574 | 0.41 |
| Cirobu.g00000825 | KH.C1.574 | MS4A12; MS4A18 | 0.79163172 | 5.08E-104 | 6.83E-100 | 0.984 | 0.787 |
| Cirobu.g00004593 | KH.C2.4 | MS4A8 | 0.98795959 | 2.50E-56 | 3.36E-52 | 0.752 | 0.347 |
| Cirobu.g00005375 | KH.C3.212 | MS4A15;MS4A18; MS4A8 | 0.39694546 | 7.83E-05 | 1 | 0.199 | 0.109 |
| Cirobu.g00005790 | KH.C3.598 | MS4A15 | 0.90634152 | 7.35E-42 | 9.87E-38 | 0.442 | 0.133 |
| Cirobu.g00010730 | KH.L114.8 | MS4A1;MS4A12;MS4A4A | 0.60792447 | 2.92E-07 | 0.00392209 | 0.333 | 0.082 |
| Cirobu.g00012421 | KH.L37.56 | MS4A12;MS4A12;MS4A15 | 0.25464204 | 3.96E-40 | 5.31E-36 | 0.248 | 0.044 |
| Cirobu.g00013381 | KH.L96.38 | MS4A4A | 0.46195901 | 0.00803779 | 1 | 0.121 | 0.036 |
| Cirobu.g00013577 | KH.S1090.3 | MS4A8 | 0.5656217 | 1.14E-31 | 1.53E-27 | 0.236 | 0.053 |

The genes are identified by unique gene ID (Cirobu.gxxxxxxx), KyotoHoya (KH) gene model ID, and Top BLASTP hit in humans. Single-cell RNAseq data from [5] using Seurat [33] include Average Log(FC), p value, adjusted p value, Pct-1, and Pct-2 according to the re-analysis of [14].

Also enriched in ACCs were 9 genes encoding members of the MS4A gene family (Table S1), which encodes receptors expressed in mammalian necklace OSNs. To confirm the specificity of expression we cloned approximately 1kb upstream of a *Ciona MS4A* gene (KH.C1.484) transcription start site and inserted it upstream of *mCherry* to form *MS4A>mCherry*. This was electroporated into *Ciona* zygotes and examined alongside ßγ-crystallin (detected with an antibody [32]) and the regulatory region of the Notch ligand *Delta-like* (KH.L50.6) [53] which we also cloned upstream of *mCherry* (forming *Delta-like>mCherry*). ßγ-crystallin and *Delta-like>mCherry* mark the ACCS and PSN respectively (Figure 6H-J). *MS4A>mCherry* expresses in the palps (Figure 6A) where it overlaps with ßγ-crystallin in the external digitiform protrusions of the ACCs, though transgene expression also includes other palp cells (Figure 6B-G).

**Figure 6.**
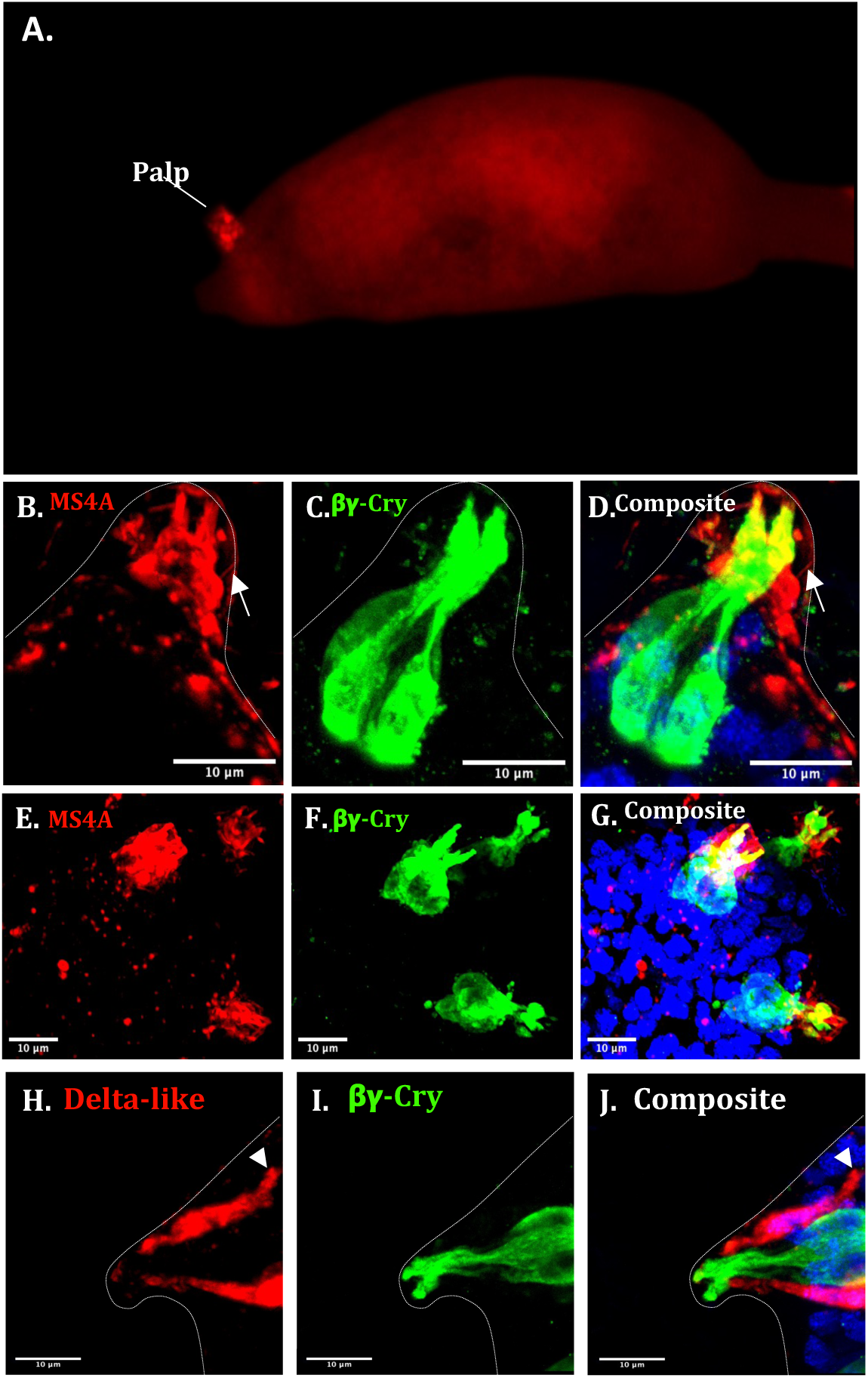
The *Ciona* MS4A chemoreceptor KH.C1.484 is expressed in the ACCs. (A) *Ciona* larva head expressing the reporter, *MS4A>mCherry*. The mCherry reporter expression is seen in the palp. A single palp is labelled here which is a common occurrence of mosaicism during transgenesis. (B-G) Confocal projections of palp cells of transgenic larvae (24 hpf) with (B-D) an individual palp and (E-G) three palps of a single animal. mCherry (labelled with an anti-mCherry antibody) from *MSA4>mCherry* is in red, ßγ-Crystallin labelled with a specific antibody (see methods) is in green labelling the ACCs, and nuclei are labelled with DAPI in blue. Note both overlap in expression (yellow in the composites in D and G) and MS4A signal that is non overlapping (in red, arrow on D). H to J show *Delta-like>mCherry* which marks the PSNs compared to ßγ-Crystallin (axon with arrowhead), with the composite showing these do not overlap. These are 2D projections of 3D reconstructions, video files of 3D projections are in Additional Files 9-11. The dotted outline marks the palp in B-D and H-J. Scale bars 10 μm.

### Molecular evolutionary features of ascidian MS4A genes

Genes encoding chemosensory receptors, including vertebrate MS4A genes, may show evidence of positive selection, and tend to be located in gene clusters and show frequent lineage specific duplications [19]. To gain further insight into *Ciona MS4A* genes we extracted MS4A sequences from the ascidian species *C. intestinalis* Type A (*Ciona robusta*), *Ciona savignyi, Botryllus schlosseri, Botryllus leachi, Halocynthia aurantium, Halocynthia roretzi, Molgula occidentalis, Molgula occulta, Phallusia fumigata and Phallusia mammalita*. A molecular phylogenetic analysis showed evidence of lineage specific expansion of *MS4A* genes in different ascidian lineages (Additional File 4). In the *Ciona intestinalis* Type A (*C. robusta*) genome there are 29 identified MS4A genes. Among those, 14 are linked to members of the orphan 7 transmembrane (TM) receptor gene family suggested to be putative chemoreceptors (Johnson et al., 2020). This chromosomal linking pattern is similar to other chordates where *MS4A* genes are found in clusters and are linked to 7 TM ORs. In *C. intestinalis* Type A (*Ciona robusta*) these comprise two clusters on chromosome 3 of 7 genes and 4 genes respectively, and one on chromosome 4 with 2 genes, with the remaining gene on chromosome 8 (Additional File 5).

Vertebrate chemosensory genes also tend to show characteristic patterns of sequence evolution, with frequent changes in sites encoding the extracellular loops of the protein, and with evidence for positive selection sometimes identifiable at these ligand-binding sites [19, 54]. Multiple sequence alignments of *Ciona* species MS4A sequences revealed substantial intra-specific (Figure 7A) and inter-specific (Figure 7B) diversity within the two extracellular domains of the proteins. To test if positive selection was involved, we sought to calculate the ratio of non-synonymous (dN) to synonymous (dS) substitutions, dN/dS (=ω), across sites in the alignment. However, despite morphological similarity, ascidian genomes are very diverged with even that between the congeneric species *Ciona intestinalis* and *Ciona savignyi* providing a very low background of sequence conservation.

**Figure 7.**
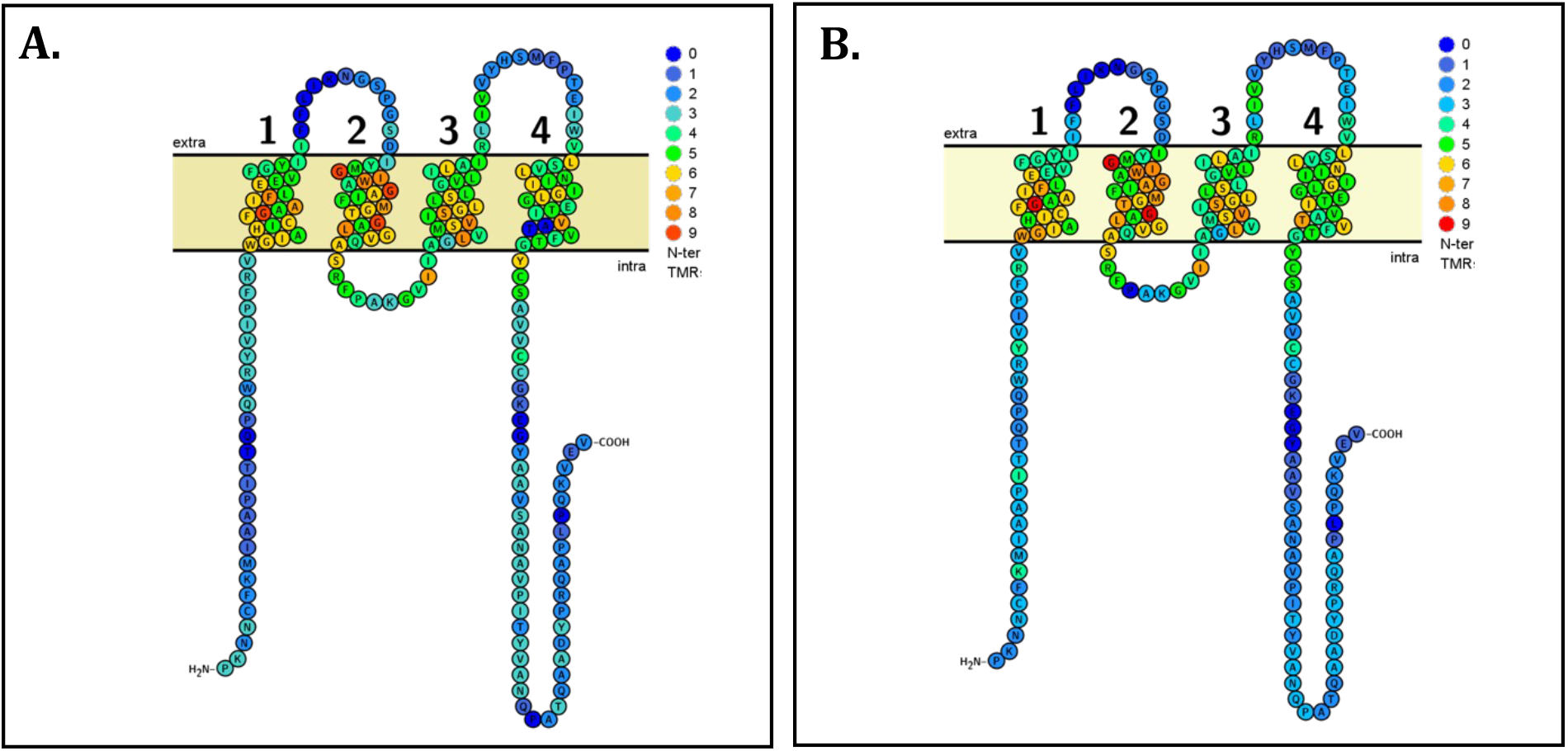
Topographical representations of *Ciona* MS4A (KH.C1.484) primary sequence with amino acid conservation heat mapped. (A, B). Residues that are more conserved are shown in warmer colours, whereas residues that are less conserved are depicted in colder colours, on a 0-9 scale. A shows an intraspecific comparison through amino acid conservation between all MS4A paralogues in *Ciona intestinalis* Type A (*Ciona robusta*) while B shows interspecific conservation between all MS4A homologs in this species and *Ciona savignyi*. The4 extracellular domains are numbered.

To circumvent this problem, we instead identified clades of recently diverging MS4A paralogues specific to individual species (Figure 8). The colonial ascidian *Botryllus schlosseri* turned out to be the most appropriate for such an analysis as the molecular phylogeny revealed the presence of clades deriving from recent paralogous expansions (Figure 8; Additional File 4). We analysed 6 clades in total, with the number of genes per clade varying from 4 to 18 and the total tree length in number of substitutions per codon from 0.62 to 9.38 (Table 2). The average ω-values across alignments (model M0) never reached 1, the threshold for inferring positive selection (Table 2).

**Table 2.** Characteristics of the codon sequence alignments per clade.

| CLADES | N | C | T | $\omega$ |
| --- | --- | --- | --- | --- |
| Clade 1 | 4 | 372 | 0.7 | 0.8141 |
| Clade 2 | 4 | 396 | 1.3 | 0.3166 |
| Clade 3 | 4 | 372 | 0.62 | 0.4874 |
| Clade 4 | 8 | 378 | 3.11 | 0.3948 |
| Clade 5 | 12 | 363 | 4.96 | 0.3812 |
| Clade 6 | 18 | 291 | 9.38 | 0.3149 |

N, number of sequences; C, number of codons; T, total tree length (in number of substitutions per codon); $\omega$ , average dN/dS over all alignment positions analysed (model M0).

**Figure 8.**
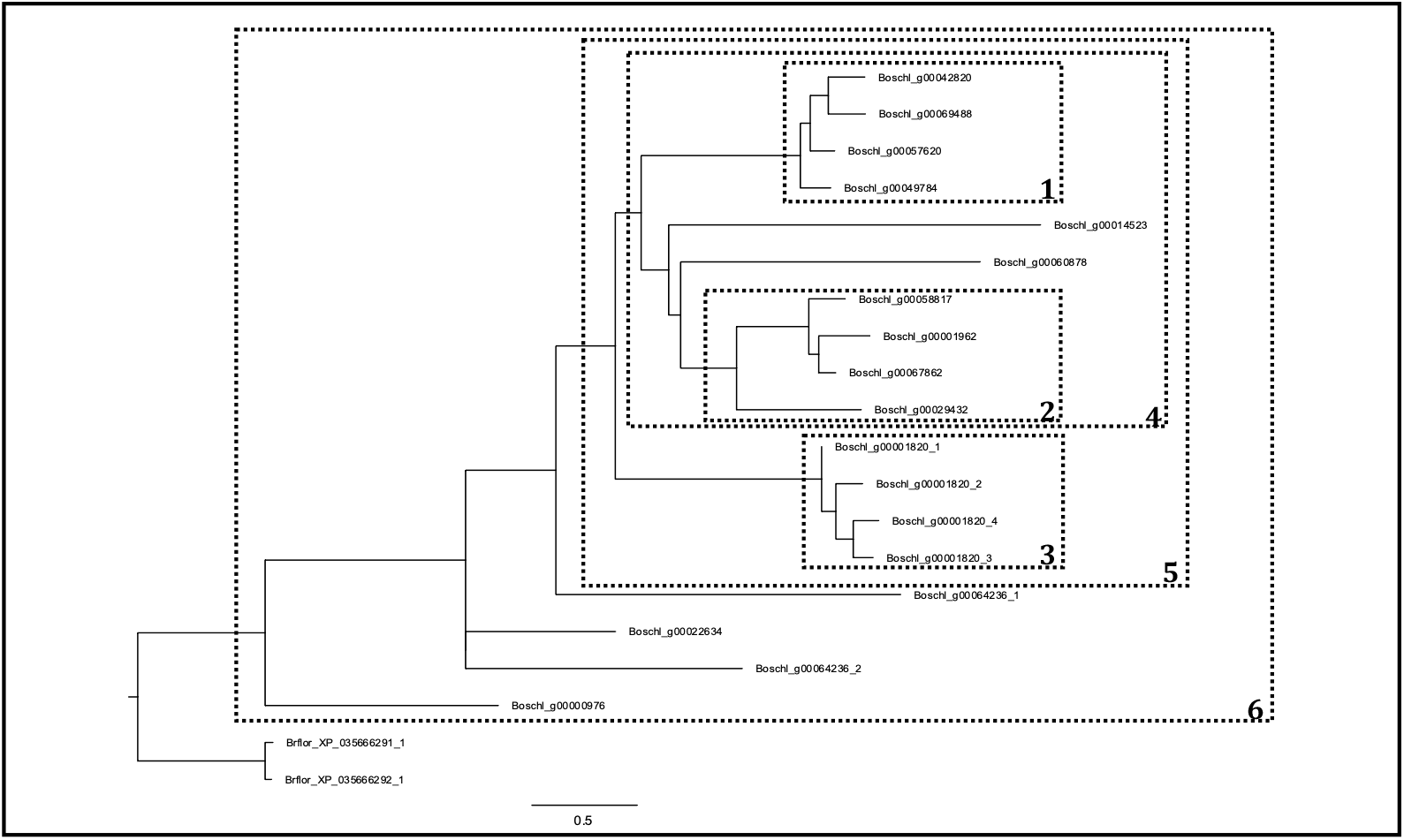
Clades of *B. schlosseri* paralogous genes analysed for evidence of positive selection. Each clade is boxed by a dashed line, and the numbering correspond to the results summarized in Table 3. All trees are rooted with amphioxus (*B. floridae*) MS4A proteins as an outgroup. The scale bars represent the number of substitutions per site.

**Figure 9.**
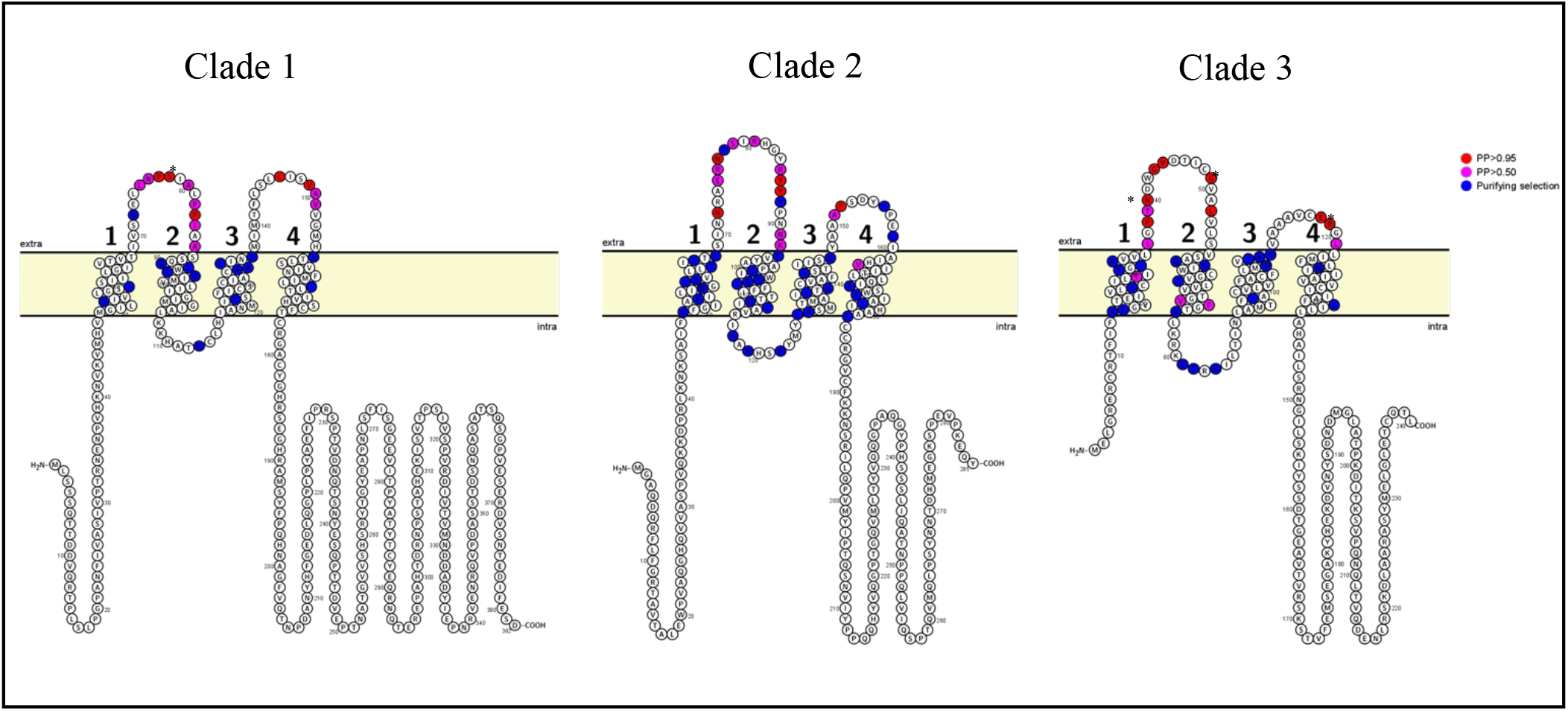
Schematic representation of the positively selected sites located on the different MS4A domains. Amino acids under high probability to be under positive selection are in magenta (PP >0.5) and red (PP>0.95). Sites with a PP>0.99 have an additional asterisk. Sites under purifying selection are in blue. The numbers indicate transmembrane domains 1 to 4.

To investigate site-specific values of ω multiple sequence alignments for each clade were fed to the codon substitution models M1a, M2a, M7 and M8 (see method for details). The presence of positively selected sites in the analyzed paralogs was supported by the highly significant likelihood ratio tests (LRT) in all clades when comparing M1a versus M2a or M7 versus M8 (Table 3). The bigger the clade the fewer positive selected sites were found, due to the increasing level of divergence between paralogues. The strongest signal of positive selection was in the most recently expanded clade (Clade 3, LRT P-values in Table 3). The Bayes empirical Bayes (BEB) method was then used to predict the location of such sites under the models M2a and M8. This identified the specific codons with a strong posterior probability (PP) to belong to the ω > 1 site class.

**Table 3.** Results of likelihood ratio tests and parameters estimates under the best-fitting model for each clade

| CLADES | M1a vs M2a<br>LRT (P-value)<sup>a</sup> | M7 vs M8<br>LRT (P-value)<sup>b</sup> | Parameters estimates<sup>c</sup> | Positive sites<sup>d</sup> |
| --- | --- | --- | --- | --- |
| Clade 1 | 0.000034549 | 0.000033956 | p0= 0.94775<br>p1= 0.05225<br>$\omega$= 27.36303 | **28 L 0.788**,29 N 0.517,30 P **0.977\***,31 S **0.994\*\***,33 A 0.609,35 P **0.685**,36 P **0.955\***,37 I 0.507,39 A 0.605,**98 L 0.945**,**101 V 0.983\***,**102 A 0.766**,103 V 0.574 |
| Clade 2 | 0.001722930 | 0.000021385 | p0= 0.84271<br>p1= 0.15729<br>$\omega$= 4.17122 | **24 S 0.987\***,27 T **0.847**,28 K **0.814**,29 R **0.957\***,31 E **0.622**,33 T **0.714**,34 Y **0.592**,35 S **0.964\***,36 Y **0.982\***,40 N **0.629**,41 V 0.533,100 G **0.905**,101 L **0.954\***,113 I 0.757 |
| Clade 3 | 0.000011343 | 0.000005698 | p0= 0.90223<br>p1= 0.09777<br>$\omega$= 29.95506 | **16 T 0.586**,25 I **0.628**,27 P **0.976\***,28 T **0.685**,29 N **0.995\*\***,30 G **0.977\***,31 V **0.960\***,36 G **0.997\*\***,39 L **0.984\***,59 V 0.526,60 F **0.824**,68 C **0.878**,101 E **0.965\***,102 R **0.997\*\***,104 L **0.928** |
| Clade 4 | 0.001946302 | 0.000001075 | p0= 0.90160<br>p1= 0.09840<br>$\omega$= 3.16466 | 26 L 0.589,27 L **0.893**,28 N **0.975\***,30 S **0.995\*\***,32 A 0.628,34 P **0.762**,35 P **0.960\***,92 T **0.876**,97 V **0.921** |
| Clade 5 | 0.000050308 | 0.000000000 | p0= 0.88319<br>p1= 0.11681<br>$\omega$= 2.48143 | **23 S 0.780**,25 E 0.612,26 N **0.999\*\***,27 P 0.823,**28 S 0.995\*\***,29 I 0.642,32 P **0.987\***,33 P **0.878**,93 V **0.990\*\*** |
| Clade 6 | 1.000000000 | 0.000023868 | p0=0.94775<br>p1= 0.05741<br>$\omega$= 2.39634 | 22 Y 0.710, **23 P 0.992\*\***, **24 G 0.915** |

<sup>a,b</sup> P-value obtained with the Likelihood Ratio Test (LRT) after M1a and M2a models comparison or M7 and M8 models comparison
<sup>c</sup>Parameters estimated under the model M8: p0 = proportion of sites that follow a beta distribution with 10 classes ( $0 < \omega < 1$ ); p1 = proportion of sites in the extra class with $\omega > 1$
<sup>d</sup>Positive site under positive selection according to M7 vs M8 models comparison with a PP > 0.50. The sites with PP > 0.95 have a single asterisk while sites with PP > 0.99 have double asterisk. The residues in bold are the one also identified with the M1a vs M2a comparison which is more stringent. The numbers and letters correspond to the positions and relevant amino acids of the sequences that were present on top of the multiple codon alignments

Table 3 presents all the positive sites with PP >0.5, however as amino acid sites are generally accepted to be under positive selection if they have at least a PP > 0.95 [42] these are further emphasized in the table. To examine how these sites might relate to protein function we mapped their location to predicted protein domains of Clade1-3 (Figure 8C). All the MS4A codons predicted to be under positive selection with PP > 0.95 are in the extracellular domains. In addition, the majority (92.5%) of sites identified as being under purifying selection are in the transmembrane domains (TM1–TM4) and the intracellular loop between TM2 and TM3.

## Discussion

### A microfluidic chip for imaging transgenic *Ciona*

Small marine larvae such as those of *Ciona* offer the prospect of linking sensory input through the network of individually identified cells and through to behavioural output. Here we report a step towards this goal with the immobilisation of live transgenic larvae in a microfluidic chip and recording of cell activity while a specific stimulus is applied. Three technical developments were needed to support this, the development of an appropriate chip and fluid control system, the delivery of a transgene to the right cells and imaging of informative fluorescence. Our chip design allows immobilisation of multiple larvae and their exposure to three different fluid streams in a controlled manner. We did find some difference in speed of onset on the left and right sides of the chip (averages of 278 msec for the left side versus 502 msec for the right side; see Methods for more details). In principle the symmetric design should mean the two sides behave the same way, so this may be due to imprecision in the mould or variation in the tubing used to supply the solutions to the lateral streams. However, speed of onset and offset on the two sides were each relatively consistent.

We used this to examine larval sensitivity to dissolved CO_2_. In total we recorded Ca^2+^ transients in 7 intact larval heads while stimulating them with CO_2_ or a pH matched control solution, and identified cells responding to these stimuli. The clearest specific response came from cells in or over the top of the sensory vesicle. Ca^2+^ levels in these cells precisely matched the onset and offset of the CO_2_ stimulus but were not triggered by the pH matched control. This shows the Ca^2+^ activity was specifically caused by the elevated CO_2_ in the test solution and not by the pH difference or by mechanosensation of the change in stream. This is the first demonstration that *Ciona* larvae can sense dissolved CO_2_ levels. This could be for predator avoidance in the plankton but might also help larvae identify an appropriate settlement site. To distinguish between these, it will be necessary to simultaneously image stimulus receipt and its response, either in the form of effector neuron activity (for example motor neurons) and/or specific behaviours such as muscle activity or triggering of metamorphosis. The cells responding to CO_2_ in this instance are most probably the aATENs based on their position in the head. However, we cannot exclude the possibility that underlying neurons within the sensory vesicle are responding to the stimulus as well as or instead of aATENs; the former is quite likely as aATENs project to this region of the sensory vesicle, so if aATENs respond we would expect neurons here to also respond [3]. Experimentally distinguishing between these possibilities will be feasible with the development of transgenic drivers specific for these cell subsets.

We also detected Ca^2+^ transients in the palps, although unlike the aATENs these responded to both stimulus and control solutions. This could reflect a response to lowered pH but could also be cells responding to mechanical force. Other experiments have demonstrated that the palps can detect mechanical stimuli and that this is part of the mechanism that initiates larval attachment to the substrate and metamorphosis [25]. It remains unclear, however, as to which cells within the palps are mechanosensory and it remains likely some cells are chemosensory as biofilm detection is part of settlement site choice. Finally, we also detected Ca^2+^ transients in the RTENs during chemical stimulation. As these cells relay sensory information from palps to sensory vesicle, their activity is most likely caused by the activity in the palps, however as we only recorded activity from this area from one embryo firm conclusions cannot currently be drawn.

Our study shows that GCaMP6m can be delivered to a defined subset of cells using transgenesis, and that its activity can be recorded to yield biological insight. Because the *DMRT* enhancer we used has quite broad anterior neural expression, we were not able to identify specific cells in our recordings, however this could be accomplished by using enhancers specific to individual cells or cell types. Some such enhancers are already and single cell sequencing data [4] will provide the route to identifying more. It is straightforward to introduce multiple transgenes simultaneously into *Ciona*, so anatomically separated cells like palp neurons, RTENs and sensory vesicle neurons could be labelled and recorded at the same time. Our methodology could also be used to study sensitivity to other potential olfactants, including those derived from biofilms from natural settlement sites or artificial structures like ship hulls. Challenges in applying this chip design remain, however. Most notably *Ciona* larvae are soft-bodied and damage is possible on loading into the chip. While cells remain alive in damaged larvae and can be recorded, connections between nerve cells may be compromised. This emphasises the importance of being able to load multiple larvae simultaneously as this allows intact larvae to be selected for imaging.

### Genomic insights into ascidian chemosensation

Previous surveys of ascidian genomes have failed to identify conclusive orthologues of vertebrate *OR* genes, or of Vomeronasal Receptor (*VR*) genes [47-52]. However ORs have been described from amphioxus [47, 55], which is more distantly related to vertebrates than is *Ciona*. ORs and VRs are members of the G-Protein Coupled Receptor (GPCR) superfamily, and divergent 7 transmembrane domain genes that may be their equivalent of GPCR ORs have been identified in *Ciona* [12, 14]. Whether these are actual chemosensors remains to be established.

Experimental confirmation that environmental CO_2_ sensing occurs in *Ciona* led us to examine the underlying mechanism of CO_2_ sensing. Based on recent description of a subset of vertebrate OSNs (the necklace OSNs) that mediate this, we asked if genes characteristic of necklace OSNs were present in *Ciona* and, if so, what could we deduce from their expression and molecular evolution. Published scRNA-seq data showed orthologues of genes involved in CO_2_ sensing in vertebrates were enriched in *Ciona* palps, specifically the ACCs. These data suggest these cells may be a site of CO_2_ sensing. They do not, however, disprove other cells as potential sensing sites, and the data are not sufficient to resolve RTEN and aATEN cell populations such that expression in these cells could be examined.

We also found 9 members of the *MS4A* receptor gene family enriched in ACCs. *Ciona intestinalis* Type A (*Ciona robusta*) has 29 members of this family in total and at least one of the remaining 9 genes (*KH. C1*.*484*) is also expressed in the ACCs, as shown by the activity of the *MS4A>mCherry* construct. This construct also drove some reporter expression in PSNs, as detected in ciliated structure typical of PSNs. We also found ascidian *MS4A* genes showed molecular evolutionary features characteristic of vertebrate *MS4A* genes in particular and of chemosensory receptors in general; clustered organisation, lineage specific gene duplication, plus elevated levels of amino acid diversity in the extracellular regions with evidence of positive selection on some of these sites. This later aspect was only identified in one species, *B. schlosseri*, which showed a sufficient number of recent gene duplications for robust analysis. Further studies including additional species would be needed to establish whether this was also the case in the *Ciona* genus.

## Conclusions

The ability to detect and discriminate chemical cues is assumed to be important for marine larvae, though this has seldom been experimentally tested. Our data establish that larvae of the ascidian *Ciona* can detect dissolved CO_2_. Dissolved gas sensation, along with other chemosensation, is important for larval behavioural ecology and may have implications for understanding biofouling and invasion biology, both significant worldwide impacts of ascidians. They also demonstrate methodology that could be used to determine if larvae can detect other chemical cues, and if so which cells are responsible and which cells downstream in the nervous system are in turn activated. In principle the methodology could be used to interrogate other aspects of neuronal circuitry as well, though this will require the development of many more cell type-specific constructs to drive GCaMP expression.

Our data also provide insight into cell type evolution in chordates. aATENs have been proposed to be homologous to vertebrate olfactory placode derived sensory cells based on their embryonic origin form the anterior neural plate border and their expression of GnRH and divergent GPCR genes [12, 14]. *Ciona* palps have been similarity proposed to be chemosensory though they are also likely mechanosensory [14, 56] and comprise at least three cell types [15]. Our data do not distinguish between mechanosensation and chemosensation as palp reporter activity responded to both stimulus and control treatments, however genomic and molecular evolution data suggest ACCs may be a site for chemosensation and homologous to CO_2_-sensitive vertebrate necklace olfactory neurons. ACCs also express GnRH [14] and we note olfactory placode derived CO_2_-sensing neurons that express GnRH have been identified in zebrafish [57]. This contrasts with the recent suggestion that ACCs may be a non-neural myoepithelial cell type homologous to vertebrate smooth muscle, myoepithelial, and cardiomyocyte cells [14]. These findings are not necessarily contradictory. It has been argued that multifunctionality is a common feature of ancient metazoan cell types [58] and ACCs could be descended from a multifunctional chemosensory cell type whose functions have been distributed to different daughter cell types during vertebrate evolution.

## Acknowledgements

We thank Dr Laurence Lemaire and Professor Mike Levine for providing the *DMRT>GFP* construct. PDMS microfluidic chip fabrication was done in the laboratory of Professor Dirk Aarts (Oxford Colloid Group, Physical and Theoretical Chemistry) under the supervision of Dr Lucia Parolini. Image acquisition was made with the kind help of Professor Mark Fricker. Thanks also go to Northney Marina for allowing animal collection. We also acknowledge the John Fell Fund and the Elizabeth Hannah Jenkinson fund for financial support.

## Availability of Data and Materials

Plasmid constructs generated in this study are available by request from the corresponding author. Raw imaging files of larval GCaMP recordings are available on figshare (https://figshare.com/articles/media/Ciona_larvae_raw_calcium_recordings/18675578). Design specifications for the microfluidic chip and chip holder are included with the manuscript as Additional Files 6 and 7.

## Competing Interests

The authors have no competing interests.

## Notes

### Competing Interest Statement

The authors have declared no competing interest.

