## Additional files 1-5 for "A microfluidic device for controlled exposure of transgenic *Ciona intestinalis* larvae to chemical stimuli demonstrates they can respond to carbon dioxide"

### SUMMARY

#### Additional Files

**Additional File 1: Figure S1 Experimental set up**
**Additional File 2: Figure S2 Chip Calibration**
**Additional File 3: Figure S3 Recordings from damaged larvae**
**Additional File 4: Figure S4 Molecular phylogenetic tree of ascidian MS4A sequences**
**Additional File 5: Figure S5 MS4A gene linkages**

**Additional data not included in submission due to file size restrictions**
Movies depicting raw recordings of all larvae range from 0.25 to 2.1Gb in size and are available on figshare:
https://figshare.com/articles/media/Ciona_larvae_raw_calcium_recordings/18675578

**Figure S1**

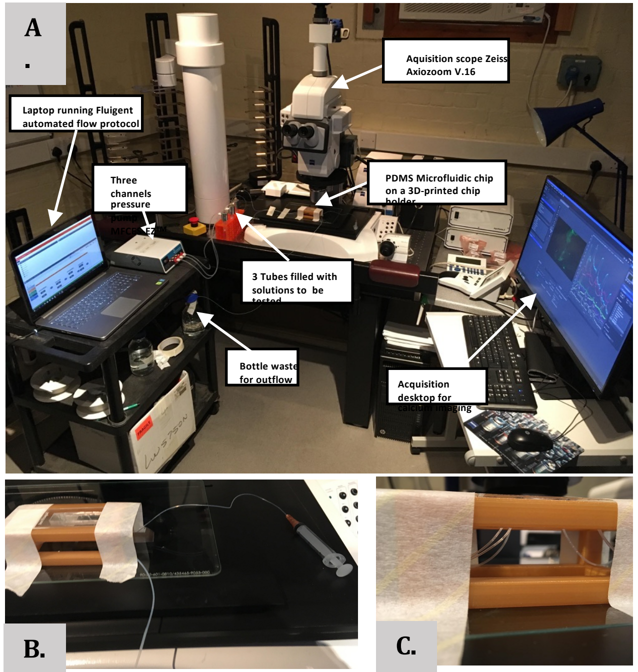


**Figure S1 Experimental set up.** (A) The controller MFCES-EZ™ with three pressure channels on the left side of a Fluorescence setero microscope Zeiss Axio zoom V.16. Each pressure channel operates a tube connected to an inlet of the chip. The bottle on the left bottom for collection of the outflow. (B) The customised 3D-printed chip holder (golden colour) holding the PDMS microfluidic chip with tape. The syringe visible on the right side of the microfluidic chip is used for animal introduction. (C) The three inlet tubings are visible on the left side while the introduction and outlet tubings are visible on the right side.

**Figure S2**

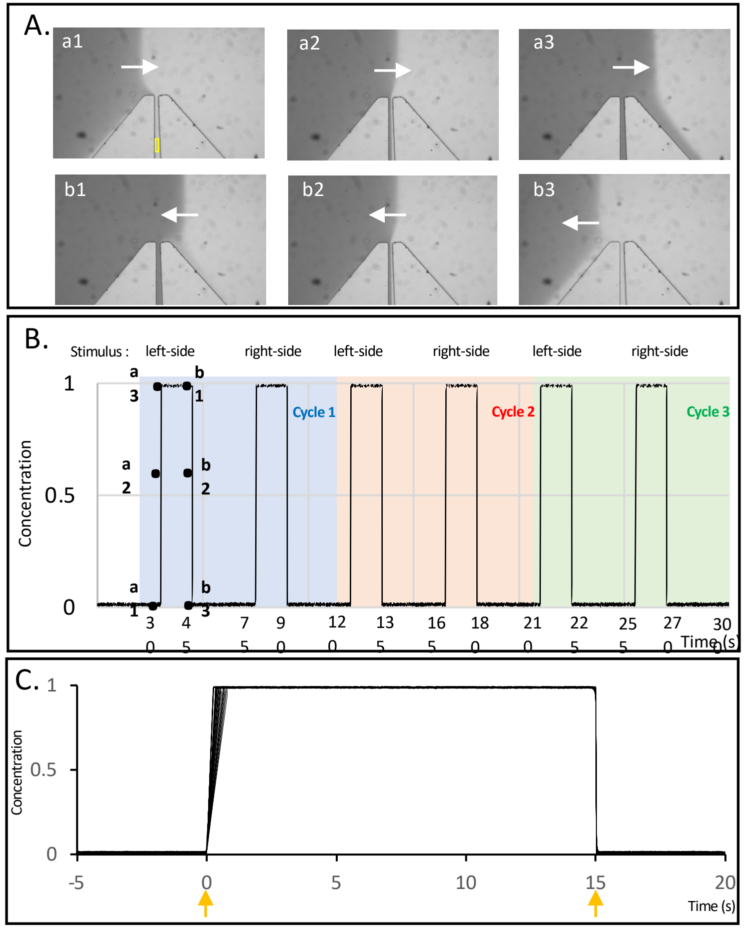


**Figure S2. Figure S2 Chip Calibration.** (A) The pictures illustrate the consecutive steps during onset ($a_1$,$a_2$,$a_3$) and offset ($b_1$,$b_2$,$b_3$) of the stimulus located on the left side of the trapping channel. The flow moves from top to bottom and the arrow indicate the horizontal movement of stream boundaries, visualised with a dye. (B) Graph representing the normalised stimulus concentration estimated from mean grey value intensity measured in a region of interest inside the trapping channel (yellow rectangle shown in a1). The graph represent the entire experimental flow protocol used during one calibration experiment. The trapping channel is exposed to either left-side stimulus or right-side stimulus during 15 sec with resting intervals of 30 sec during three repetitive cycles (blue, red, green). **.** (B) Graph representing all 60 measurements of onset/offset with the different cycles and stimulus sides together. The normalised stimulus concentration (from 0 to 1) is estimated from mean grey value intensity. The orange arrow indicate the time point when the pressure controller switches pressure rates starting at time 0 and ending at time 15.

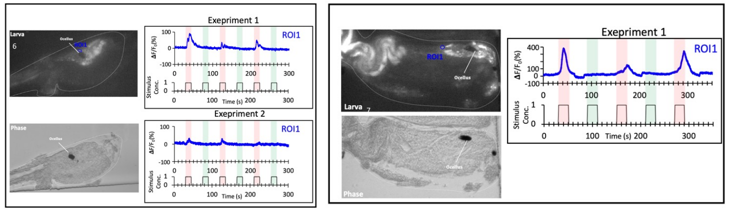


**Figure S3 Recordings from damaged larvae.** Larva 6 and 7 sustained obvious tissue damage upon introduction in the chip but showed nonetheless obvious responsiveness to the microfluidic system. The graphs show the temporal patterns of fluorescence intensity for each ROI. The red trace corresponds to exposure to the left side stimulus, i.e., the solution with high $CO_2$. The green trace is the exposure to the right-side stimulus, i.e., the flow and pH control.

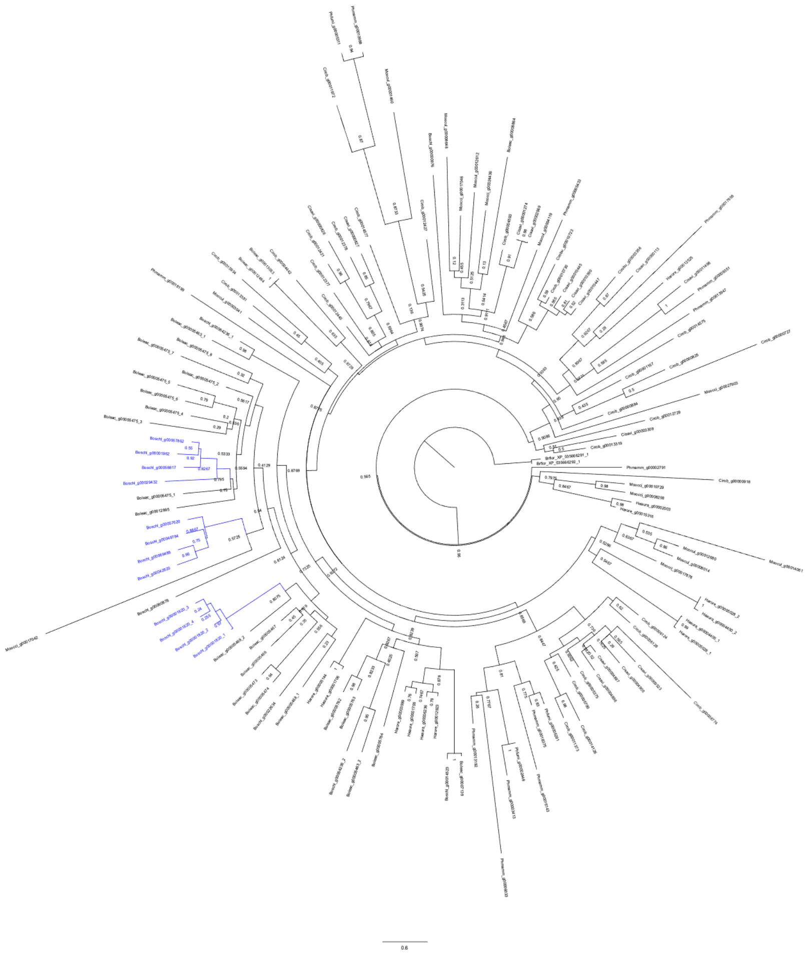

**Figure S4 Molecular phylogenetic tree of ascidian MS4A sequences.**
Circular maximum-likelihood phylogenetic tree of 128 ascidian MS4A proteins. Specific MS4a gene expansions of *B. schlosseri* are in blue. the rest of ascidian gene names and branches are in black.

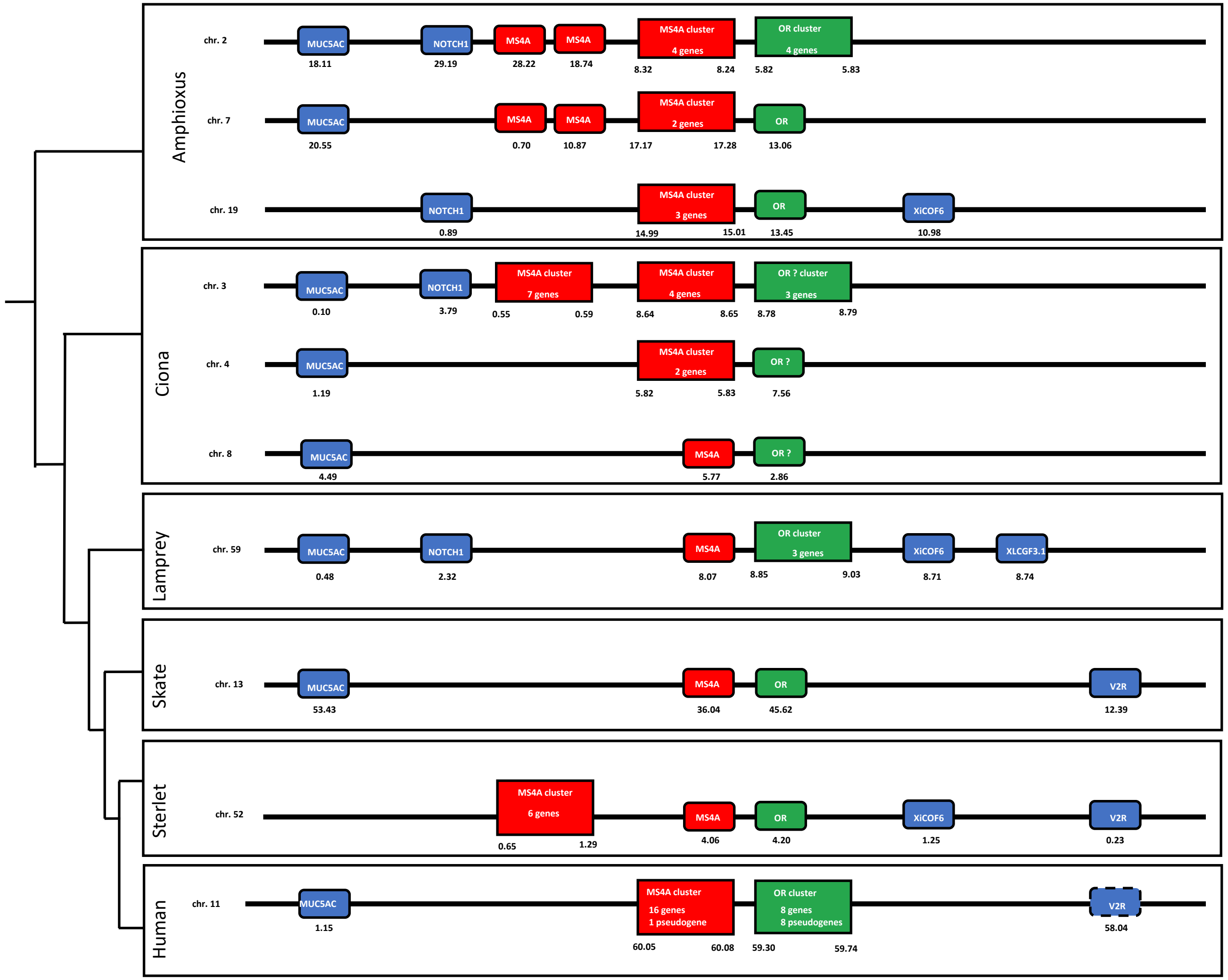


**Figure S5. Figure S5 MS4A gene linkage.** The neighbouring genes are aligned for ease of comapraison, but the distance (shown below in Mb) and order may vary. *MS4A* genes are in red, colour coding of other genes is as follows: Green, *OR* genes . Blue colour, genes linked to MS4A-OR genes on the same chromosome and their orthologues and/or paralogues in other species. Dotted outlines represent pseudogene.
